# Exploring genetic resistance to Infectious Salmon Anaemia Virus in Atlantic salmon by genome-wide association and RNA sequencing

**DOI:** 10.1101/2020.09.08.287052

**Authors:** O. Gervais, A. Barria, A. Papadopoulou, R. Gratacap, B. Hillestad, A.E. Tinch, S.A.M. Martin, D Robledo, R.D. Houston

## Abstract

Infectious Salmonid Anaemia Virus (ISAV) causes a notifiable disease that poses a large threat for Atlantic salmon breeders and producers worldwide. There is no fully effective treatment or vaccine, and therefore selective breeding to increase resistance to ISAV in commercial strains of Atlantic salmon is a promising avenue for disease prevention. Genomic selection and potentially genome editing can be applied to enhance host resistance, and these approaches benefit from improved knowledge of the genetic and functional basis of the target trait. The aim of this study was to characterise the genetic architecture of resistance to ISAV in a commercial Atlantic salmon population and study its underlying functional genomic basis using RNA Sequencing. A total of 2,833 Atlantic salmon parr belonging to 194 families were exposed to ISAV in a cohabitation challenge in which cumulative mortality reached 63% over 55 days. A total of 1,353 animals were genotyped using a 55K SNP array, and the estimate of heritability for the trait of binary survival was 0.33 (±0.04). A genome-wide association analysis confirmed that resistance to ISAV was a polygenic trait, albeit a genomic region in chromosome 13 was significantly associated with resistance and explained 3% of the genetic variance. RNA sequencing of the heart of 16 infected (7 and 14 days post infection) and 8 control fish highlighted 4,927 and 2,437 differentially expressed genes at 7 and 14 days post infection respectively. The complement and coagulation pathway was down-regulated, while several metabolic pathways were up-regulated in infected fish compared to controls. The interferon pathway was mildly activated at 7 days and showed no sign of up-regulation at 14 days post infection, implying a crosstalk between host and virus. Comparison of the transcriptomic response of fish with high and low breeding values for resistance (4 high resistance and 4 low resistance animals per time point) highlighted TRIM25 as being up-regulated in resistant fish, suggesting it may be a key antiviral gene involved in the functional genetic basis of resistance to ISAV.

## 1. INTRODUCTION

The demand for high-quality animal protein for human diets has increased steadily during the last decades and is expected to accelerate over the next thirty years, in parallel to human population growth [1]. When paired with the challenges of climate change and increased competition for land use [2], a sustainable increase in farmed animal protein production efficiency is required to meet the global food security challenge [3,4]. Aquaculture is typically resource-efficient, with high rates of feed efficiency and protein retention compared to terrestrial livestock [5], and is expected to play a major role feeding the world in the coming years. While aquaculture production has risen steadily in the recent decades [6], it can also be high-risk, in part due to infectious diseases, which pose major threats to entire production systems, with downstream impacts on efficiency and sustainability.

One such disease threat for farmed Atlantic salmon (*Salmo salar*) is infectious salmon anaemia (ISA), caused by an aquatic orthomyxovirus of the same name (ISAV). ISA is classified as a list II disease by the EU fish health directive and as a notifiable disease by the World Organisation for Animal Health (OIE), which means that entire stocks have to be culled upon detection of the virus to avoid the spread to nearby farms. While outbreaks were first detected in Norway, ISA has been observed in all major salmon producing countries [7–13]. Just two years after its first detection in Chile in 2007, ISA caused the collapse of the salmon aquaculture industry, reducing Atlantic salmon production by ~75% in two consecutive years [14]. While ISA outbreaks have been recorded almost exclusively in marine-phase farmed Atlantic salmon, the virus has also been detected in other salmonid species such as rainbow trout (*Oncorhynchus mykiss*), coho salmon (*Oncorhynchus kisutch*) and brown trout (*Salmo trutta L*) [14]. The most characteristic clinical sign of the disease is severe anaemia, often accompanied by lack of appetite and lethargic behaviour [14]. In production settings, a severe ISA outbreak can cause mortalities of above 90% [15].

ISAV is an enveloped negative-sense single stranded RNA virus member of the family *Orthomyxoviridae*, and therefore closely related to influenza viruses. Viruses of this family share similar strategies of infection, using haemagglutinin activity to enter the cells and fusion activity to escape the lysosome, followed by viral RNA replication in the nucleus of the host cell and modulation of host immune responses [16–18]. The genome of ISAV is divided in 8 segments that encode at least 10 different proteins, and the virus can be divided in two groups, the low virulence ISAV-HPR0 and the virulent ISAV-HPRΔ, which has a deletion in the highly polymorphic region of the haemagglutinin-esterase gene [19]. Currently there are no effective treatments against ISAV, and available vaccines are typically only partially protective [20]. Furthermore, vaccinated fish can still carry the virus, and can therefore become asymptomatic carriers spreading the disease to other fish. Moreover, vaccination of fish can result in positive results in ISAV detection tests, which can exclude facilities from ISA free compartment status, preventing export of fish from these facilities [21]. For these reasons, there are restriction on the use of ISA vaccines, which are actually not allowed in some countries, and other strategies are necessary to limit the impact of ISAV.

The use of genetic and genomic technologies is becoming an integral part of efforts to reduce the frequency and severity of disease outbreaks in aquaculture species [22]. Genomic selection exploits both between and within family genetic variation to improve the innate resistance of aquaculture stocks via selective breeding, with cumulative benefits every generation [21]. Several studies have shown that host resistance to ISAV has a significant additive genetic component in Atlantic salmon, with heritability estimates ranging from 0.13 to 0.40 [23–27]. Furthermore, studies using molecular markers to investigate the genetic architecture underlying this heritability have revealed putative QTL on chromosomes 3, 4, 15 and 21 [28,29], and a comparative genomic analysis highlighted potential underlying genes [30]. Several studies have also examined the host response to ISAV by profiling gene expression in tissues and cell lines [31–36]. Generally, these studies have reported a notable up-regulation of innate immunity which did not confer complete protection from the impact of the virus, and which was less marked in vaccinated or secondary-infected fish. Notably, salmon immune responses to ISAV have been reported to be tissue-dependant and tightly regulated by viral transcription [36].

Genetic improvement by selective breeding is limited by the existing additive genetic variation for the trait of interest in the population, and the ability to efficiently measure the trait, which limits the accuracy of selection and therefore genetic gain. The detection of functional genes and variants controlling disease resistance, as well as a better understanding of the genomic mechanisms underpinning disease resistance, can contribute to improve the efficiency of aquaculture breeding programmes by improvement of genomic selection methods [22]. Furthermore, this information can feed into genome editing efforts to enhance disease resistance, whether it is exploiting existing genetic variation or generating de novo mutations based on the functional basis of disease resistance [37,38]. One route to achieving this is to integrate transcriptomic data with genetic mapping data to identifying putative functional genes and pathways connected to resistance, and this approach has been applied for genetic resistance to viral and parasitic diseases in Atlantic salmon [39,40].

To assess the potential for selection of ISAV resistance in a commercial Atlantic salmon population and gain insight into the functional genetic basis of the trait, a large scale ISAV disease challenge in 2,833 Atlantic salmon parr belonging to 194 families of the SalmoBreed and StofnFiskur strains was performed. A total of 1,353 fish were genotyped for 55K SNP markers, and RNA sequencing was performed on subsets of the challenged population with divergent breeding values for resistance. These datasets were then used to: i) evaluate the heritability of resistance to ISAV in a commercial Atlantic salmon population, ii) assess the genetic architecture of the trait using a genome-wide association study (GWAS), and iii) compare the transcriptomic responses to infection and if this response varied between resistant and susceptible animals.

## 2. METHODS

### 2.1. Disease challenge and sampling

The population used for the ISAV challenge experiment comprised 2,833 parr fish (mean 37.5 g) from 194 nuclear families originating from Benchmark Genetics breeding programme. The challenge experiment and sampling were conducted in the facilities of VESO Vikan (Norway). All fish were PIT-tagged and transferred to one 4 m^3^ tank where they were acclimated for three weeks in fresh water at the following approximate conditions: temperature 12°C, stocking density 40 kg / m^3^, flow 5-6 mg O_2_ / L and photoperiod regime L:D = 24:0. Post acclimation, 300 carrier fish (Atlantic salmon from the same population) used for the cohabitation challenge were intraperitoneally injected with 0.1 mL of ISAV (Glaesvær, 080411, grown in ASK-cells, 2 passage, estimated titre 10^6^ PFU / mL) and introduced to the challenge tank with the naive fish. Mortalities were registered daily, and the trial was terminated when the mortality level dropped to baseline levels (i.e. near zero). Adipose fin tissue samples from all fish were collected and stored in ethanol for DNA extraction and genotyping. In addition, 30 of the challenged fish were terminated at each of three time points (pre-infection, 7 days post infection and 14 days post infection) for sampling of tissues for transcriptomic analyses. In addition to fin clips, the hearts of a subset of animals were collected into TRI Reagent (Sigma, UK) and stored at −20°C until RNA extraction.

### 2.2. Genetic parameter estimation

Resistance to ISAV was measured as binary survival (BS), mortalities were recorded as 0 and survivors as 1. All challenged fish were used to estimate the genetic parameters for resistance to ISAV, using a probit link function and the ASREML software v4.1 [41]. The univariate animal model used was:

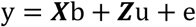

Where ***y*** is the vector of phenotypic records; ***b*** is the vector of fixed effects, which includes sex and the first two principal components as fixed effect, and body weight at PIT-tagging as covariate; ***u*** is the vector of random animal genetic effects which assumes the following normal distribution 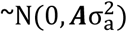, where ***A*** is the additive relationship matrix and 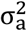 is the additive genetic variance; ***e*** is the vector of residual effects with a normal distribution assumed as 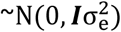, where ***I*** is the incidence matrix and 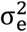 is the residual variance; and ***X*** and ***Z*** are design matrices for fixed and random effects, respectively. Heritability was estimated as:

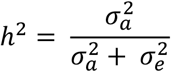

Where 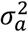 is the additive genetic variance and 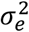 is the estimated residual variance, which was set as 1 (Gilmour et al. 2009).

### 2.3. Genotyping and GWAS

A total of 1,367 samples were successfully genotyped for a 55K Affymetrix Axiom SNP array used routinely by Benchmark Genetics in their commercial breeding programme. DNA extraction from fin clips and SNP array genotyping was performed by IdentiGEN (Dublin, Ireland). Quality control (QC) was performed using PLINK software v1.90 [42]. SNPs with minor allele frequency (MAF) lower than 0.05 or significantly deviating from Hardy–Weinberg Equilibrium (HWE) (p < 1e-6) were removed for further analyses; SNPs and individuals with a call rate lower than 99% were also excluded.

A weighted single-step GBLUP approach (wssGBLUP) was used to estimate genomic heritability and to identify the genomic regions associated with ISAV resistance. This approach takes into account fish with phenotypic and genotypic data, but also considers those fish with phenotypes and no genotypes, combining data from all challenged fish (connecting genotyped and not genotyped fish through the pedigree) [43]. Pedigree and genotypic information are combined to create an ***H*** matrix [44]. Thus, the inverse of this ***H*** matrix is:

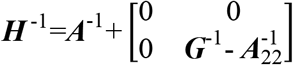

Where ***A***^−1^ is the inverse of the pedigree-based relationship matrix, 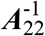 represents the inverse of the ***A*** matrix, but only considering the genotyped fish, and ***G***^−1^ is the inverse of the genomic relationship matrix. The statistical model for the genetic parameter estimation and for the genome-wide association study is identical as the one mentioned above, but replacing the ***A*** matrix by the **H** matrix. Genomic heritability using the **H** matrix was estimated as described above. The variance obtained on the first iteration of wssGBLUP for each SNP (single-step GWAS) was used as the weight in the analyses. The variances were estimated based on the allele frequency and marker effect [45]. A threshold model was fitted for BS using the THRGIBSS1F90 function of BLUPF90 [46], and a total of 200,000 Markov Chain Monte Carlo (MCMC) iterations were fitted. From these, 20,000 were burned-in, and 1 from every 50 of the remaining 180,000 samples were saved.

Additionally, a mixed linear model, using the leaving-one-chromosome-out (LOCO) approach was fitted to identify SNPs associated with resistance to ISAV, through the GCTA v.1.92.2. software [47]. The fitted model was identical as the one described for the wssGBLUP approach, although a G matrix was used. For a SNP to be significantly associated at genome-wide level with resistance to ISAV, it must surpass the Bonferroni-corrected significance threshold (a/n), where a and n represent the significance level (0.05) and the number of SNP that surpassed the QC, respectively.

Finally, the p-values for each SNP and the proportion of the genetic variance explained by 20 adjacent SNP window were plotted with R/CMplot.

The wssGBLUP approach was used for the estimation of the genetic parameters and the genome-wide association study was also used to predict the genomic estimated breeding values (gEBVs) for resistance to ISAV, estimating the genetic resistance and susceptibility of the fish sampled for transcriptomic experiments.

### 2.4. RNA extraction and RNA sequencing

For each timepoint (control, 7 and 14 days post infection) 4 fish with high breeding values for resistance and 4 fish with low breeding values for resistance from 8 different families were selected according to their estimated genomic breeding value for ISAV resistance. Heart RNA was extracted from preserved tissue samples (n= 24) in TRI Reagent (Sigma, UK) and RNA extracted following the manufacturer’s instructions. The RNA pellet was eluted in 15 μL of nuclease-free water and quantified on a Nanodrop 1000 spectrophotometer (NanoDrop Technologies) prior to DNAse treatment with QuantiTect^®^ Reverse Transcription kit (Qiagen). The quality of the RNA was examined by electrophoresis on a 1% agarose gel (Sigma Aldrich), prepared in Tris-Acetate-EDTA (TAE) buffer, stained with 0.05% ethidium bromide (Sigma Aldrich) and run at 80 V for 30 min. Sample concentration was measured with Invitrogen Qubit 3.0 Fluorometer using the Qubit RNA HS Assay Kit (ThermoFisher Scientific). PolyA RNA-Seq libraries were prepared using Illumina’s TruSeq RNA Library Prep Kit v2 by Oxford Genomic Centre, and sequenced on an Illumina Novaseq6000 as 150bp paired-end reads yielding an average of 51M reads per sample (minimum 38M).

### 2.5. RNA-Seq analyses

Raw reads were quality trimmed using Trimgalore v0.6.3. Briefly, adapter sequences were removed, low quality bases were filtered (Phred score < 20) and reads with less than 20 bp were discarded. Trimmed reads were then pseudoaligned against the Atlantic salmon reference transcriptome (ICSASG_v2 Annotation Release 100; Lien et al., 2016) using kallisto v0.44.0 [49]. Transcript level expression was imported into R v3.6 [50] and summarised to the gene level using the R/tximport v1.10.1 [51]. Differential expression analysis was performed using R/Deseq2 v1.22.2 [52], and genes with False Discovery Rate adjusted p-values < 0.05 were considered to be differentially expressed. Kyoto Encyclopedia of Genes and Genomes (KEGG) enrichment analyses were carried out using KOBAS v3.0.3 [53]. Briefly, salmon genes were annotated against the KEGG protein database [54] to determine KEGG Orthology (KO). KEGG enrichment for differentially expressed gene lists was tested by comparison to the whole set of expressed genes (average of > 10 normalised reads) in the corresponding tissue using Fisher’s Exact Test. KEGG pathways with ≥ 5 DE genes assigned and showing a Benjamini-Hochberg FDR corrected p-value < 0.05 were considered enriched.

### 2.6. Ethics statement

The challenge experiment was performed at VESO Vikan with approval from the Norwegian Food Safety Authority, National Assignments Department, approval no. 16421, in accordance with the Norwegian Animal Welfare Act.

### 2.7. Data availability

RNA sequencing raw reads have been deposited in the NCBI’s Short Read Archive (SRS) repository with accession number PRJNA647285.

## 3. RESULTS

### 3.1. Disease challenge and genetic parameters of ISAV resistance

The ISAV cohabitation challenge on 2,833 fish belonging to 194 families from Benchmark Genetics commercial breeding programme showed substantial variation in mortality rate between families (Fig. 1A), with values ranging from 7 to 100 %, suggesting the presence of a genetic component underlying resistance to ISAV in this population. Mortalities began at day 19 and reached 63 %, with most mortalities occurring between day 22 and 28 (Fig. 1B). The pedigree-based heritability for resistance to ISAV was estimated to be 0.13 ± 0.05.

**Fig. 1.**
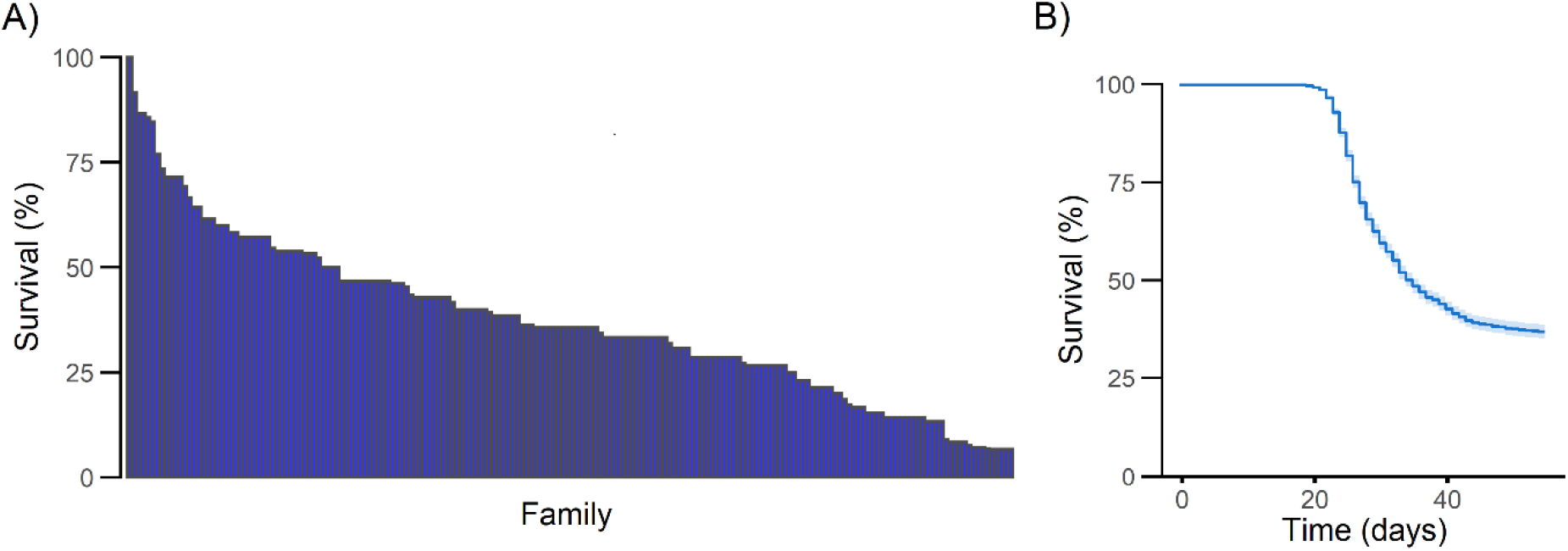
Patterns of mortality observed during the ISAV challenge. A) The percentage of survival for each full-sibling family at the end of the challenge, and B) The total percentage of survival in the population throughout the duration of the challenge.

### 3.2. Genetic architecture of ISAV resistance

A subset of the challenged population (n = 1,353) was genotyped using a 55K SNP array. After QC processing, a total of 43,346 SNPs and 1,103 fish remained for downstream analyses. Genomic heritability estimated using the weighted single-step GBLUP model was 0.33 ± 0.04, which is notably higher than the pedigree estimate. The single SNP genome-wide association analysis revealed a significant QTL in chromosome 13 (Fig. 2A; a single SNP in chromosome 16 also reached the significance threshold, but was not supported by other SNPs in the region). The QTL in chromosome Ssa2 explained ~3% of the genetic variance in resistance to ISAV, while five other genomic regions each explained more than 1%, with the largest-effect detected in Ssa18 (4.8%) (Fig. 2B, Supplementary Table 1). Overall, the data supported a polygenic basis to host resistance to ISAV, with minor effect loci distributed across several chromosomes.

**Fig. 2.**
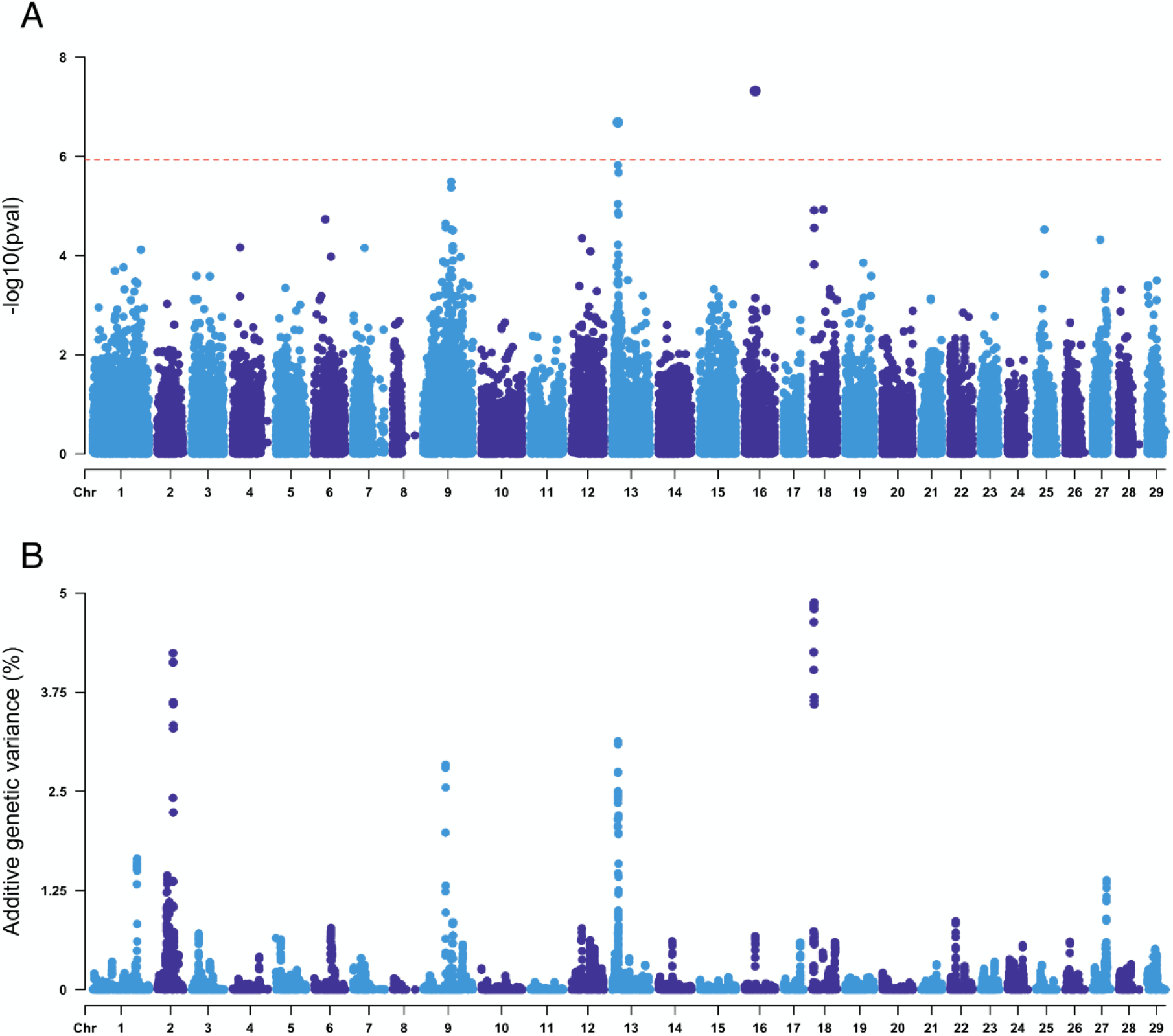
Weighted single-step genome-wide association analyses for resistance to ISAV in the challenged Atlantic salmon population. A) shows the p-value for each SNP in a single SNP GWAS, and the red dotted horizontal line represents the significance threshold (p-value < 0.05 after Bonferroni correction); B) shows the percentage of additive variation explained by windows of 20 consecutive SNPs. 17 SNPs are placed in scaffolds not assigned to chromosomes (ICSASG_v2; Lien et al. 2016) and are not shown. These unassigned SNPs explained less than 0.01 % of the genetic variance and were not significantly associated with resistance to ISA.

### 3.3. Transcriptomic response to ISAV

Based on the genomic estimated breeding values for resistance to ISAV and family mortalities, 4 resistant and 4 susceptible animals were selected at each of three timepoints (0, 7 and 14 days post infection). The average GEBVs of resistance to ISAV for the more resistant and more susceptible groups across all timepoints were 0.05 and 0.41 (survival = 0, mortality = 1), respectively, with average family survival rates of 64 and 17 % for each group. The transcriptome of the heart samples from these animals was sequenced using Illumina technology, obtaining an average of 51 million of reads per sample. Principal component analyses highlighted that control and infected samples clustered separately according to the two first principal components, which explained 20 and 13 % of the total variance (Fig. 3). However, there was no clear differentiation between the challenged timepoints, nor between the resistant and susceptible samples (Supplementary Figure 1).

**Fig. 3.**
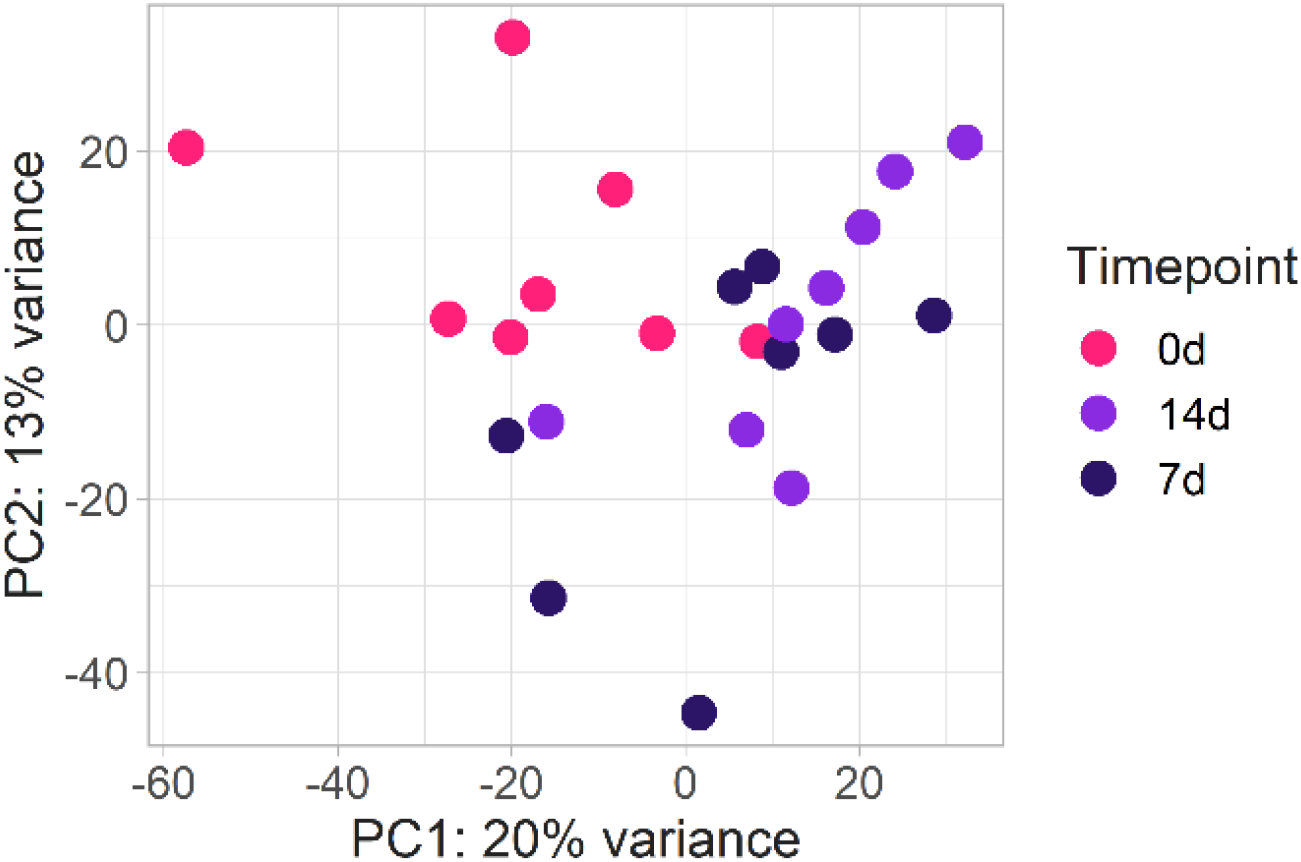
Principal Components Analysis showing the clustering of the heart RNA-Seq data.

Although the differentiation between the sampling timepoints based on all transcript data was not clear cut, there was a notable response to ISAV observed in the heart samples both at 7 and 14 days post infection, with 4,927 and 2,437 differentially expressed genes when compared to controls, respectively (Fig. 4 & Supplementary Table 2). Most of the genes differentially expressed at 7 days were also differentially expressed at 14 days post infection (1511 genes; Fig. 4A). Several genes in the interferon pathway are mildly up-regulated at 7 days post infection (Fig. 4B), suggesting an activation of the antiviral pathway early in infection, as shown by several interferon regulatory factor (IRF) mRNAs being significantly higher in infected samples compared to controls (albeit with small fold change). However, at 14 days post infection, there was little evidence for differential expression of interferon genes, with the exception of one copy of Interferon-induced Very Large GTPase 1 (GVINP1, another copy of the gene is down-regulated) (Fig. 4C). It is also noteworthy that two genes encoding Mx, a key antiviral gene, showed significantly lower expression at 14 dpi. Other well-characterised immune genes showed differential expression, such as Tumor Necrosis Factor alpha, up-regulated at 7 days but not at 14 days post infection. Down-regulation of numerous complement genes was observed at both timepoints, and in fact KEGG pathway enrichment analyses (Table 1 & Supplementary Table 3) revealed a clear and increasing down-regulation of the complement and coagulation cascades pathway during infection. Almost all the complement and coagulation cascade genes showing putative downregulation at 7 dpi showed even larger negative fold changes in expression at 14 dpi, and additional genes from the same pathways showed statistically significant down-regulation (Supplementary Table 4). Similarly, a consistent up-regulation of numerous metabolic processes is observed at both 7 and 14 dpi. Interestingly, several of the pathways typically activated during innate immune response to viruses, such as interferon, interleukin or inflammation pathways, are not enriched amongst the set of up- or down-regulated genes, although the pathway HTLV-I (human T-lymphotropic virus type 1) infection is down-regulated at 7 dpi, and so are certain signalling pathways closely related to innate response such as FoxO and mTOR signalling.

**Fig.4.**
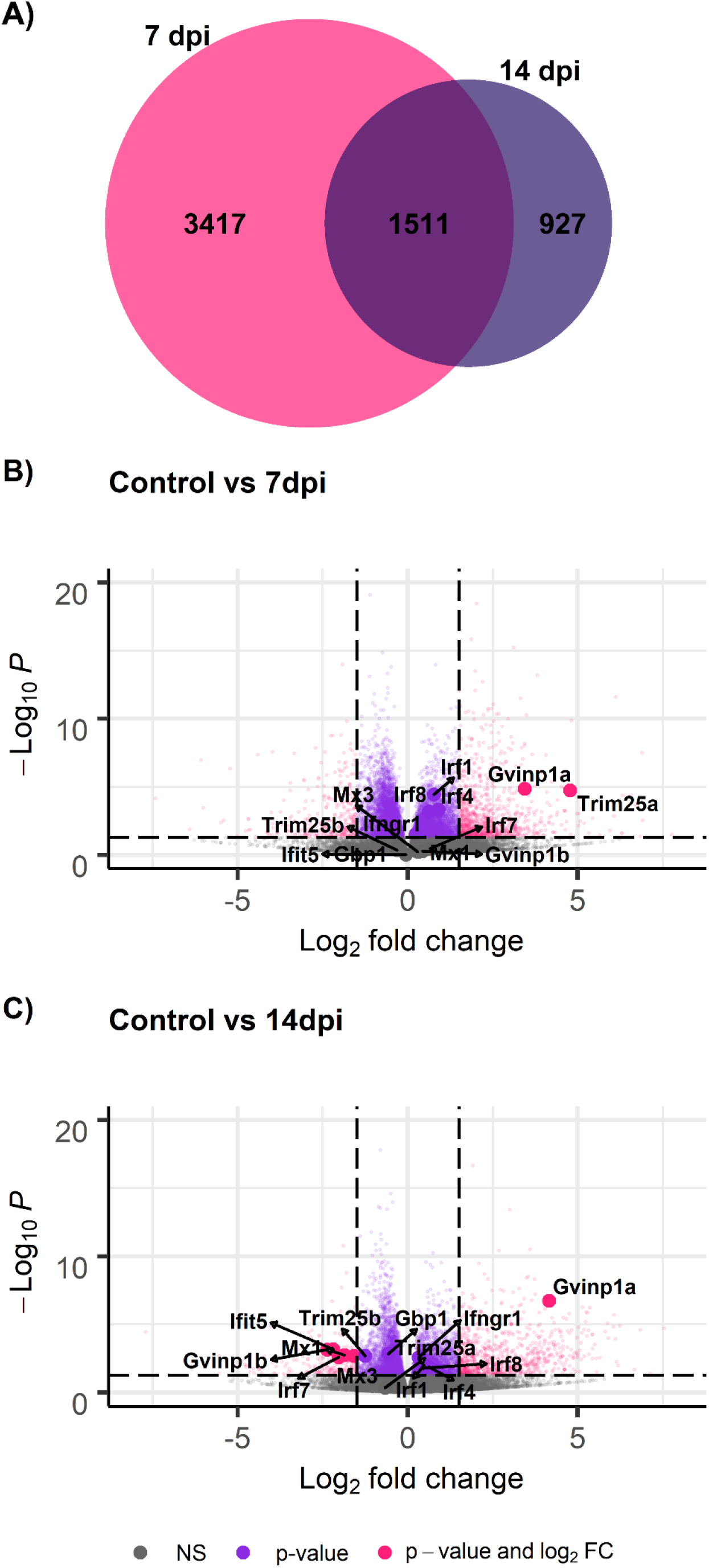
Differential expression of transcripts between ISAV-infected and control fish. **A)** Venn diagram depicting the number of common and unique genes showing differential expression at 7 and 14 days compared to control. B) Volcano plot showing the differential expression and differentially expressed interferon genes in control vs 7 dpi, and C) controls vs 14 dpi. Each point in the plots represents a gene, with its log_2_ fold change in the x-axis and its −log_10_ p-value in the y-axis. Genes are classified in 4 categories depending on their FC and FDR corrected p-value: i) grey = p-value > 0.05 and log_2_ fold change between −1.5 and 1.5; ii) yellow = p-value > 0.05 and log_2_ fold change < −1.5 or > 1.5; iii) light blue = p-value < 0.05 and log_2_ fold change between −1.5 and 1.5; and iv) purple = p-value < 0.05 and log_2_ fold change <−1.5 or > 1.5).

**Table 1.** Selected KEGG pathways identified as enriched for differential expressed genes.

| **7 dpi** |  |  |  |  |  |  |  |
| --- | --- | --- | --- | --- | --- | --- | --- |
| **Up-regulated** |  |  |  | **Down-regulated** |  |  |  |
| **KEGG** | **N** | **FE** | **p** | **KEGG** | **N** | **FE** | **p** |
| Carbon metabolism | 83 | 6.01 | $10^{-16}$ | Complement and coagulation cascades | 23 | 2.73 | 0.002 |
| Aminoacyl-tRNA biosynthesis | 39 | 2.98 | $10^{-13}$ | FoxO signalling pathway | 36 | 1.90 | 0.010 |
| Citrate cycle (TCA cycle) | 38 | 2.27 | $10^{-11}$ | HTLV-I infection | 50 | 1.67 | 0.012 |
| **14 dpi** |  |  |  |  |  |  |  |
| **Up-regulated** |  |  |  | **Down-regulated** |  |  |  |
| **KEGG** | **N** | **FE** | **p** | **KEGG** | **N** | **FE** | **p** |
| Carbon metabolism | 75 | 9.31 | $10^{-30}$ | Complement and coagulation cascades | 57 | 12.76 | $10^{-36}$ |
| Glycolysis / gluconeogenesis | 31 | 4.50 | $10^{-11}$ | Staphylococcus aureus infection | 24 | 5.31 | $10^{-15}$ |
| Biosynthesis of amino acids | 35 | 6.99 | $10^{-10}$ | Systemic lupus erythematosus | 17 | 7.17 | $10^{-5}$ |

KEGG, KEGG pathway; N, number of genes differentially expressed assigned to the corresponding KEGG pathway; FE, fold enrichment; p, false discovery rate corrected p-value.

### 3.4. Genomic signatures of resistance to ISAV

To assess the functional genomic basis of resistance, 4 fish of high resistance breeding values and 4 fish of low resistance breeding values were compared at each of the three timepoints (pre-challenge, 7 and 14 days post infection). There were a relatively small number of significantly differentially expressed genes between resistant and susceptible fish (13-18 DEG per timepoint; Fig. 5 & Supplementary file 5), but they included several innate immune response genes such as E3 ubiquitin/ISG15 ligase TRIM25 (involved in innate immune defence against viruses; more expressed in resistant 7 dpi, logFC = 3.90), interferon-induced very large GTPase 1 (more expressed in resistant 14 dpi, logFC = 1.31), or transcription factor Kruppel-like factor 2 (regulates inflammatory processes; less expressed in resistant controls, logFC = −1.03) (Fig. 5).

**Fig. 5.**
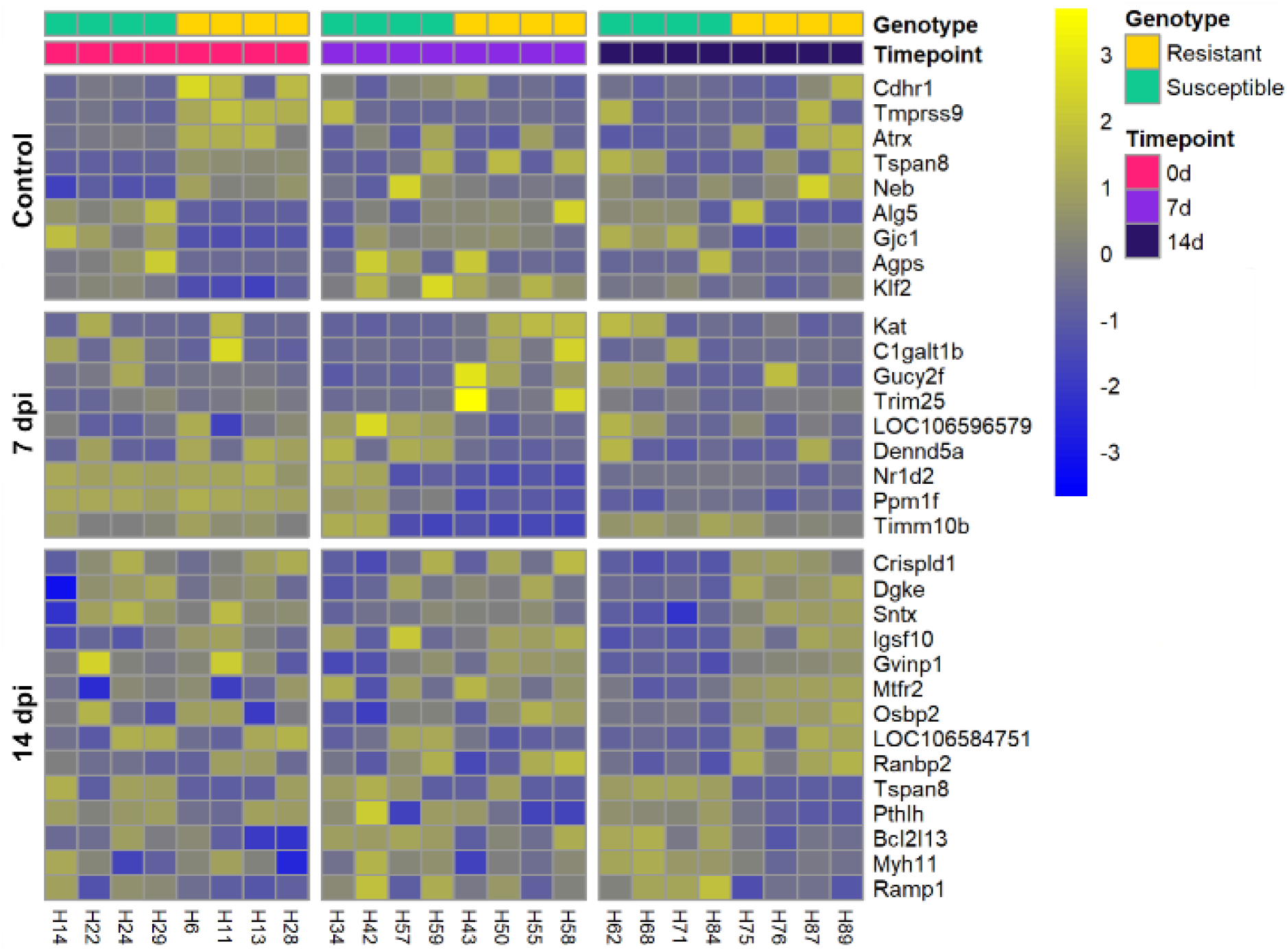
Heatmap showing the patterns of expression of genes differentially expressed between resistant and susceptible fish in each sample at all of the three timepoints.

## 4. DISCUSSION

In this study, the genetic and genomic basis of resistance to ISAV in Atlantic salmon was characterised in a large population of Atlantic salmon parr derived from 194 families of a commercial breeding programme. The trait of binary survival (reflecting host resistance) shows a moderate genetic component and is therefore amenable to selection; heritability of resistance was estimated at 0.13 and 0.33 with the pedigree and genomic relationship matrices, respectively. Higher heritability estimates using genomic relationships have been observed compared to pedigree-based estimates in previous studies investigating disease resistance in aquaculture species [55,56], potentially due to high linkage disequilibrium due to recent selective breeding causing overestimation of additive genetic variance using genomic markers [56]. Nonetheless, these results are in line with the range of previous heritability estimates for resistance to ISAV in Atlantic salmon (0.13-0.40; [23–27]). The genetic architecture of resistance to ISAV in this population was polygenic, and major QTL were not observed, as is common for disease resistance traits [56–61]. Yet, four genomic regions explaining > 2.5% of the genetic variation in resistance to ISAV were detected. Two previous studies found resistance QTL for ISAV on chromosomes *Ssa03*, *Ssa04*, *Ssa15* and *Ssa25* [28,29], however, in our study, no signal of resistance was detected in these chromosomes; this is not surprising considering the different origins of the populations and the polygenic nature of resistance to ISAV (and the different nature of the challenge, through intraperitoneal injection, in [29]).

The only genomic region with a convincing and significant association in the single-SNP GWAS approach with resistance to ISAV was found in chromosome 13, explaining ~3% of the genetic variance. Within 50 Kb of the most significant SNPs is the DDB1- and CUL4-associated factor (DCAF1; GeneID 106566667). This protein is a substrate-specific adaptor of the DDB1-CUL4-X-box E3 ubiquitin ligase complex, which mediates proteasome-mediated degradation and plays key roles in several processes, including T-cell and B-cell development [62]. Importantly, this protein is directly targeted by HIV-I and HIV-II in humans to hijack the E3 ubiquitin ligase complex to allow infection and replication [63,64]. Although not significant, a genomic region in chromosome 18 contained SNP windows which explained the most genetic variance (~5%). One of the most significant SNPs in *Ssa18* (5,781,403 bp) is located in the DNA mismatch repair protein Mlh1 gene (GeneID 106576838), which was reported to be involved in influenza A infection in human cell cultures, specifically contributing to cell survival [65]. Mlh1 is part of the MMR pathway, which prevents the accumulation of oxidative DNA lesions contributing to non-lytic viral clearance and enhanced cell survival [65]. While these genes are interesting candidates, we found no evidence of their functional involvement in the response to the virus; nonetheless, these genes could show tissue-specific responses or not depend on expression changes to have an impact on the course of the disease.

### Putative virus-induced immunomodulation of transcriptional response to ISAV in salmon

A relatively large transcriptomic response to ISAV infection was observed in the heart of Atlantic salmon at both 7 and 14 days post infection. This response was characterised by a down-regulation of the complement and coagulation cascades; surprising considering the frequently observed symptoms of the disease (erythrophagia and haemorrhages; [9]). In addition, a lower number of genes showed differential expression at 14 dpi than at 7 dpi, but the number of differentially expressed genes involved in the complement and coagulation cascades increased, suggesting potentially progressive immunomodulation by ISAV. Orthomyxoviruses are capable of subverting the host complement system through different mechanisms [66]. A general up-regulation of various metabolic pathways was also observed both at 7 and 14 dpi. Widespread metabolism dysregulation is commonly observed in diseased fish [67–69], although generally down-regulated and ascribed to a physiological response of the host to infection to adjust cellular homeostasis. Nonetheless, pathogens also reprogram the cellular metabolism of infected cells to favour their replication [70], and therefore the observed dysregulation could be a consequence of the host-virus interaction. In fact, Influenza infection strategy includes mechanisms to alter host transcription and translation [71].

The observed interferon response was weaker than expected, with a few interferon-related genes showing higher expression levels at 7 days than in controls, and typically downregulation at 14 days post infection. While Mx2 had higher expression levels than controls at 14 days, there was no change in the expression of Mx1, which has been show to confer resistance to ISAV in chinook salmon cells [72]). In general, the gene expression patterns suggest that the innate immune response was relatively mild at 7 dpi and 14 dpi. This differs from previous studies on ISAV-infected salmon [32,33,35,36]. There are two possible, non-exclusive explanations for this result. First, tissue-specific regulation seems to play an important role during ISAV infection [36], and previous research has been focused in immune-tissues, which are more likely to exhibit dysregulation of genes associated with immunity. Secondly, a previous study on the response of Atlantic salmon heart to ISAV only reported significant differences in the expression of immune genes at 21 days post infection, albeit some genes started to show an upward trend at 13 days [34]. Therefore, the relatively early timepoints studied here, in comparison to other studies, might explain the weak innate immune response. However, we do observe a clear transcriptomic response, which considering the stronger response to ISAV at 7 dpi than at 14 dpi might suggest a mechanism of immunoevasion of the virus, mainly affecting the interferon pathway (a clear up-regulation of complement genes is still observed 14 days post infection). This immunoevasion might not be effective later in the infection when viral levels are high, which is consistent with the reported positive correlation between virus abundance and the magnitude of the immune response [33,36,73]).

The down-regulation of genes involved in the interferon response (i.e. mx, irf7 or ifit5) in infected samples at 14 days post infection (and lack of up-regulation of other interferon genes at this time point) suggests that ISAV modulates this process as part of its infection and replication strategy. Previous studies have demonstrated that segments 7 and 8 of the ISAV genome produce proteins with antagonist IFN properties, binding directly to IRFs [74] and down-regulating type I IFN transcription activity [75]. Segment 7 specifically inhibits the transcription of mx [76], which has been shown to confer resistance to ISAV in chinook salmon cells [72]. This active suppression of the interferon system is consistent with previous studies in Influenza in human and chicken [77,78]. Additionally, SOCs and NLRc3 genes, which limit the inflammatory response [79,80], are up-regulated in infected samples, and cytokine induction is not clearly observed (especially at 14 days post infection), supporting the theory of the negative regulation of the innate immune response during ISAV infection.

### TRIM25 regulation could be involved in resistance to ISAV in Atlantic salmon

Although the overall antiviral response was less striking than expected, certain immune genes are up-regulated in response to ISAV (TLRs, gvinp1, TRIM25). Importantly, both gvinp1 and TRIM25, interferon stimulated genes (ISG), were found to have higher average expression in resistant than in susceptible samples. Gvinp1 is a protein directly induced by the interferon pathway, but its function is not fully understood [81,82]. On the other hand, the function of the E3 ubiquitin / ISGA15 ligase TRIM25 is well-characterised in mammals, where TRIM25 is responsible for the ubiquitination of RIG-I, leading to the activation of the downstream pathway and increased interferon production [83], crucial for antiviral innate immunity. Interestingly, Influenza A virus non-structural protein 1 (NS1) specifically inhibits TRIM25-mediated ubiquitination of RIG-I [84]. TRIM25 can also inhibit viral RNA synthesis through direct binding to the viral RNA polymerase complex (independent of its ubiquitin ligase activity), an activity that can also be inhibited by the viral NS1 [85]. In summary, TRIM25 plays a vital role in the host response to Influenza infection in mammals, and is actively modulated by Orthomixoviruses. However, it is unclear whether this function is conserved in teleost, because although fish TRIM genes show features suggesting a role in innate immunity, they show important clade-specific diversifications [86]. Nonetheless, many TRIM genes are induced upon viral infection, and some have been shown to trigger antiviral activity in vitro [87]. In common carp, a gene also annotated as TRIM25 was identified as a promising candidate for Koi herpesvirus resistance [88], which suggests its relevance in antiviral responses is well conserved. TRIM25 up-regulation in ISAV resistant samples suggests that it is a good target for future functional studies, and genome editing to change the sequence or expression of this gene could increase the resistance of Atlantic salmon to ISAV.

## CONCLUSION

Resistance to ISAV is moderately heritable and shows a polygenic architecture amenable to genome-assisted selection schemes, albeit a significant QTL was discovered in chromosome 13 explaining around 3% of the genetic variance and could be prioritised prior validation in follow-up studies. The heart transcriptomes of selected genetically resistant and susceptible samples revealed a complex response, which suggests a host-pathogen crosstalk regulating the innate immune response and more specifically the interferon pathway. In line with its polygenic architecture, the transcriptomic signatures of resistance are diverse in nature; nonetheless, TRIM25 stands out as a particularly relevant candidate for further functional studies on resistance to ISAV. Genome editing experiments should inform the role of this gene in the progression of the disease, and help obtain salmon stocks with increased resistance to ISAV in the future, leading to increased stability, food security and fish welfare.

## Funding sources

The authors gratefully acknowledge funding from BBSRC (BB/R008612/1, BB/R008973/1), in addition to BBSRC Institute Strategic Programme Grants to the Roslin Institute (BB/P013759/1 and BB/P013740/1).

## Competing interests

A commercial organisation (*Benchmark Holdings plc*) was involved in the development of this study. BH and AET work for Benchmark. The remaining authors declare that they have no competing interests.

